# Leveraging HILIC/ERLIC Separations for Online Nanoscale LC-MS/MS Analysis of Phosphopeptide Isoforms from RNA Polymerase II C-terminal Domain

**DOI:** 10.1101/2024.10.08.617299

**Authors:** Scott B. Ficarro, Deepash Kothiwal, Hyun Jin Bae, Isidoro Tavares, Gabriela Giordano, Stephen Buratowski, Jarrod A. Marto

**Affiliations:** Department of Cancer Biology and Blais Proteomics Center, Dana-Farber Cancer Institute, Boston, MA 02215, U.S.A.; Department of Pathology, Brigham and Women’s Hospital and Harvard Medical School, Boston, MA 02115, U.S.A.; Department of Biological Chemistry and Molecular Pharmacology, Harvard Medical School, Boston, MA 02115, USA; Department of Oncologic Pathology, Dana-Farber Cancer Institute, Boston, MA 02215, U.S.A.

**Keywords:** ERLIC, HILIC, Phosphopeptide, RNA Polymerase II C-terminal Domain, Mass Spectrometry

## Abstract

The eukaryotic RNA polymerase II (Pol II) multi-protein complex transcribes mRNA and coordinates several steps of co-transcriptional mRNA processing and chromatin modification. The largest Pol II subunit, Rpb1, has a C-terminal domain (CTD) comprising dozens of repeated heptad sequences (Tyr1-Ser2-Pro3-Thr4-Ser5-Pro6-Ser7), each containing five phospho-accepting amino acids. The CTD heptads are dynamically phosphorylated, creating specific patterns correlated with steps of transcription initiation, elongation, and termination. This CTD phosphorylation ‘code’ choreographs dynamic recruitment of important co-regulatory proteins during gene transcription. Genetic tools were used to engineer protease cleavage sites across the CTD (msCTD), creating tryptic peptides with unique sequences amenable to mass spectrometry analysis. However, phosphorylation isoforms within each msCTD sequence are difficult to resolve by standard reversed phase chromatography typically used for LC-MS/MS applications. Here, we use a panel of synthetic CTD phosphopeptides to explore the potential of hydrophilic interaction and electrostatic repulsion hydrophilic interaction (HILIC and ERLIC) chromatography as alternatives to reversed phase separation for CTD phosphopeptide analysis. Our results demonstrate that ERLIC provides improved performance for separation of singly- and doubly-phosphorylated CTD peptides for sequence analysis by LC-MS/MS. Analysis of native yeast msCTD confirms that phosphorylation on Ser5 and Ser2 represents the major endogenous phosphoisoforms. We expect this methodology will be especially useful in the investigation of pathways where multiple protein phosphorylation events converge in close proximity.

---

Advances in technologies for phosphoproteomics, including enrichment methods [1-6], automation [7-11], chromatographic separation[12-14], mass spectrometer data acquisition and analysis schemes [13, 15-21], along with bioinformatic tools [22-27], enable systematic and quantitative analysis of cell signaling. Use of phosphoproteomic approaches in ‘discovery’ or ‘screen’ mode can provide a detailed activity map for cellular response to a myriad of endogenous or exogenous stimuli. Researchers have used these data to illuminate disease-specific pathways [28-30], decipher response to clinical drugs [31-33], and identify modifications on key effectors that regulate cellular processes [34, 35]. Despite this progress it remains difficult to decipher mechanism or interrogate cellular processes when key effectors are decorated with multiple proximal modifications or contain functional domains comprising highly phosphorylated, degenerate sequences. In these cases, new analytical approaches are required to deconvolute the contribution of individual modifications to cellular function.

Eukaryotic RNA polymerase II (RNA pol II) transcribes all mRNAs and many non-coding RNAs (ncRNAs). This essential and highly conserved protein complex carries a unique C-terminal domain (CTD) on its largest subunit, Rpb1 (**Figure 1A**). The yeast CTD consists of 26 tandem repeats of the heptad sequence Tyr1-Ser2-Pro3-Thr4-Ser5-Pro6-Ser7. Metazoans can have up to 52 repeats, which often include many non-consensus heptamers. Each heptad contains five amino acid residues that can be differentially phosphorylated, with individual sites dynamically regulated during RNA pol II progression from initiation to elongation to termination (**Figure 1A**). The resulting “CTD code” creates specific binding sites for co-regulatory proteins involved in transcription, RNA processing, and chromatin modification that function at specific stages of transcription [36-38]. For example, phosphorylation at Ser5 correlates with initiation and early elongation, helping recruit mRNA capping enzymes and the histone H3K4 methyltransferase Set1. Ser2 phosphorylation predominates in later transcription to couple mRNA polyadenylation and H3K36 methylation. Additional roles have been proposed for phosphorylations at Tyr1, Thr4, and Ser7 [36-38].

**Figure 1.**
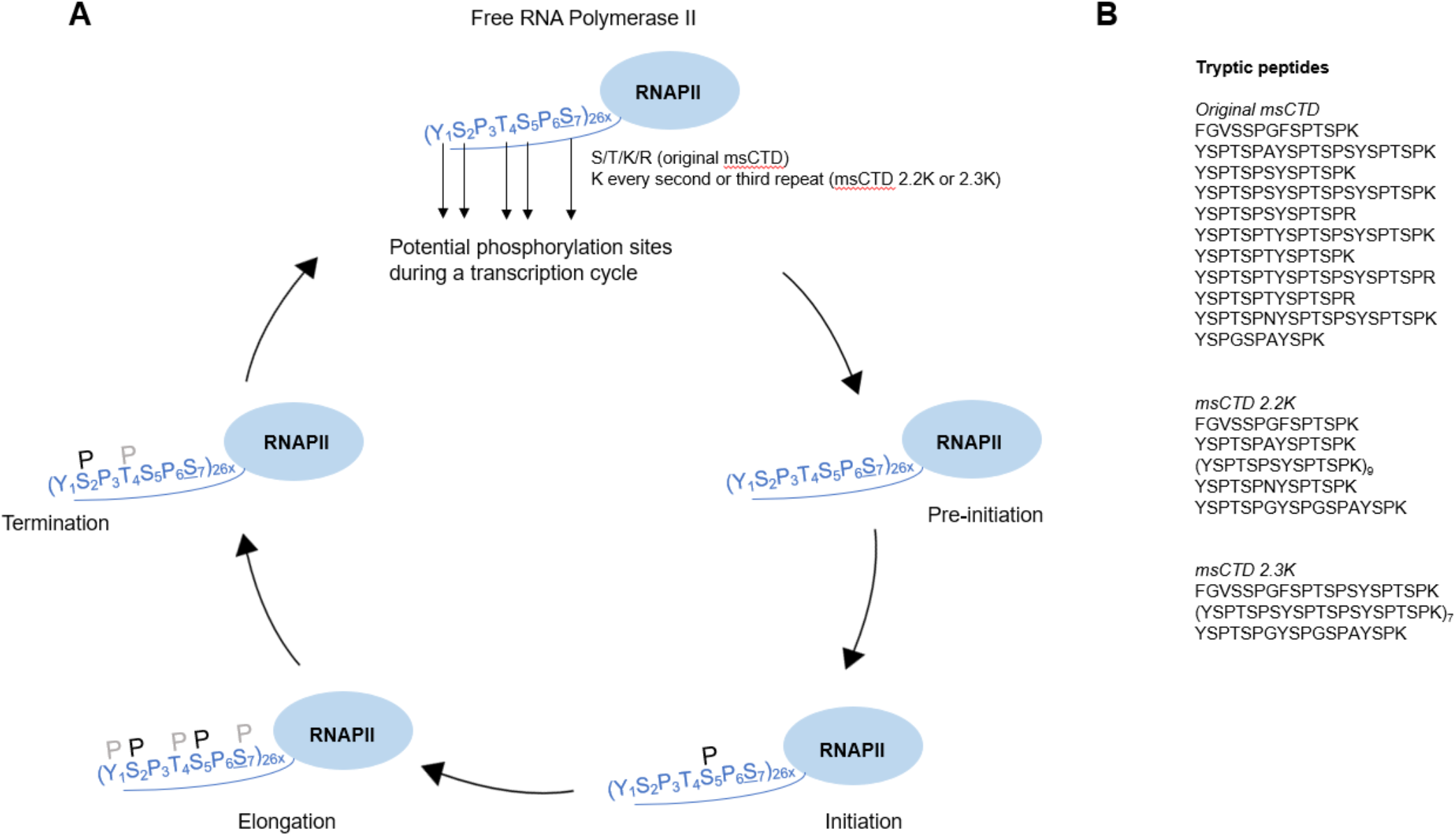
The CTD code and engineered variants of msCTD designed for (A) positional resolution and (B) co-addition of degenerate CTD peptide sequences for amplified signal accumulation in mass spectrometry. (A) CTD phosphorylation cycle. The C-terminal domain (CTD) of RNA pol II consists of tandem repeats of the heptad sequence Tyr1-Ser2-Pro3-Thr4-Ser5-Pro6-Ser7. Each heptad contains five phosphorylatable amino acid residues that are dynamically phosphorylated during different phases of transcription initiation, elongation, and termination (indicated by “P”). (B) Peptides generated by trypsin digestion of original msCTD, msCTD 2.2K and msCTD 2.3K. “Original msCTD” provides for positional resolution of phosphorylation across the entire CTD, while 2.2K and 2.3K produce degenerate peptides that concentrate signal in 9 (2.2K) or 7 (2.3K) tryptic peptides.

Current models of CTD phosphorylation are based almost exclusively on the use of phospho-specific antibodies. While these are excellent tools in general, antibodies have certain limitations with respect to quantifying discrete phosphorylation sites on the CTD. Antibody epitopes usually span several amino acids. Because 5 of the 7 amino acids that comprise the CTD are subject to phosphorylation, attempts to interrogate a specific Ser, Thr, or Tyr may be confounded by phosphorylation on one or more neighboring residues. In addition, quantitation with antibodies is problematic, as both the CTD and antibodies are multivalent, leading to non-linear signals. Furthermore, spatial information is lost as antibodies cannot distinguish phosphorylation at a given site across the numerous heptad repeats.

To overcome these issues, we previously modified yeast Rpb1 to create an “msCTD” amenable to mass spectrometry analysis(**Figure 1B**). Yeast CTD contains no basic residues, so a limited number of lysine and arginine residues were strategically incorporated to produce appropriately sized tryptic fragments. In cases of redundant peptides, a few Ser7 positions were changed to threonine, a very commonly observed substitution over evolution. This approach maintained functional polymerase activity and yielded sequence-unique CTD peptides after trypsin digestion [39]. A similar approach was independently reported in mammalian cells [40].

Using msCTD, we mapped heptad-repeat specific phosphorylation patterns from different stages of transcription, as well as changes resulting from mutation of kinases and phosphatases. High levels of phosphorylation at Ser5 and Ser2 were detected, with behavior confirming expectations from the CTD code model. No differences were apparent between phosphorylation at proximal and distal repeats. However, phosphorylations of Tyr1, Thr4, and Ser7 were nearly two orders of magnitude lower than Ser2 and Ser5 [39], which was difficult to reconcile with proposed functions for these modifications [36-38]; however, previous efforts to characterize phosphorylation at these sites was largely based on use of antibodies which, as noted above, are subject to a various confounding effects.

Although our genetic approach successfully allowed MS interrogation of tryptic CTD peptides, we subsequently discovered that phosphoisoforms of individual msCTD heptads co-eluted under standard reversed phase chromatography conditions typically used in LC-MS/MS. The resulting chimeric MS/MS spectra may complicate accurate localization of the phosphate moieties within the heptad sequence. For example, less abundant phosphoisoforms could have been obscured by stronger signals from Ser2 and Ser5 phosphorylation.

Here we use a panel of chemically synthesized CTD phosphopeptides to show that standard, low-pH reversed phase (RP) chromatography cleanly separated Tyr1P CTD peptides, but the serine/threonine phospho-isomers eluted in a single, unresolved peak. These observations motivated us to explore new LC-MS/MS-compatible chromatographic methods that can provide improved resolution of msCTD phosphopeptides. Although high-pH RP and hydrophilic interaction chromatography (HILIC) increased resolution of some msCTD species, we determined that electrostatic repulsion hydrophilic interaction chromatography (ERLIC) was superior for resolving these phosphopeptide isoforms. ERLIC buffers suitable for online LC-MS/MS detection of singly and doubly phosphorylated CTD peptides were characterized. We expect that our ERLIC approach will provide a powerful method to decipher CTD phosphorylation patterns, as well as other proteins regulated by combinatorial phosphorylation on neighboring residues.

## EXPERIMENTAL SECTION

Phosphorylated CTD synthetic peptides (>95% purity) were synthesized by Tufts university core facility or JPT Peptide Technologies. The HILIC column (TSK-Gel, 5 µm 200Å 4.6 × 250 mm) was purchased from Tosoh. ERLIC column and guard column were purchased from PolyLC (Polywax LP 5 µm 300Å 4.6 × 200 mm with Javelin guard column 20 × 4 mm µm 300Å). ZIC-HILIC column (Sequant, 5 µm 200Å 4.6 × 250 mm) was purchased from Sigma Aldrich. XBridge C18 column (XBridge, 5 µm 300Å 4.6 × 250 mm) was from Waters (Milford, MA). LC grade water, isopropanol, and acetonitrile were from Fisher Scientific. Peptides were resolved on an Agilent 1200 HPLC system with quarternary pump, autosampler, column compartment, and UV detector. Detailed gradient conditions are given in Figure Legends and Supplementary Information.

Online ZIC-HILIC and ERLIC were performed with a self-packed (15 cm of 5 µm, 300Å PolyWAXLP or 5 µm ZIC-HILIC) column with integrated emitter tip. Peptides were injected with a CTC-PAL autosampler and gradient eluted (NanoAcquity UHPLC, Waters, Milford, MA) into a 5600 Triple TOF mass spectrometer (AB-Sciex, Framingham, MA). TOF MS spectra were acquired m/z 300-2000. High sensitivity MS/MS spectra were acquired with unit resolution *m/z* 100-1800.

RNA pol II carrying the Rpb1-msCTD2.2K subunit was isolated from yeast strain YSB3609 (MATa, ura3Δ0, leu2 Δ0, trp1Δ::hisG, his3Δ1, rpb::NatMX, Rpb3-TAP::HIS3 [pCK859-msCTD2.2K]) using tandem affinity purification, followed by msCTD purifiction as previously described [39]. Recombinant msCTD2.2K was expressed in E. coli as a GST fusion using plasmid pGEX-4T-3-msCTD2.2K and purified using standard methods. Proteins were diluted in 100 mM ammonium bicarbonate, reduced (10 mM DTT, 56 °C, 30 minutes), alkylated (22.5 mM iodoacetamide, room temperature, 30 minutes, protected from light), and digested with trypsin overnight at 37 °C. Peptides were desalted by SP3 beads as described[41]. Eluates were dried by vacuum centrifugation and reconstituted in 80% acetonitrile 0.1% TFA for ERLIC. Peak areas were determined using mzStudio software[42].

## RESULTS AND DISCUSSION

Profiling phosphorylation of the RNA pol II CTD by proteomics is challenging due to the lack of suitable protease sites across heptads as well as sequence repetition and the potential of phosphorylation on 5 of the 7 amino acids comprising each heptad. Even after engineering tryptic residues (msCTD) to create sequence-unique peptides (**Figure 1**), the presence of multiple phosphorylation sites led to problematic co-elution of phosphoisomers during LC-MS/MS (**Supplementary Figure 1**). This situation confounds efforts to quantify and localize phosphorylation sites. To facilitate rapid evaluation of different conditions for improved chromatographic separation we used synthetic diheptad phosphopeptides (**Table 1**). For clarity, we use 3 letter amino acid abbreviations (ex. Ser 5) to describe heptad positions, whereas single letters (ex. pS12) designate which residue is phosphorylated within the diheptad peptide sequence.

**Table 1.**
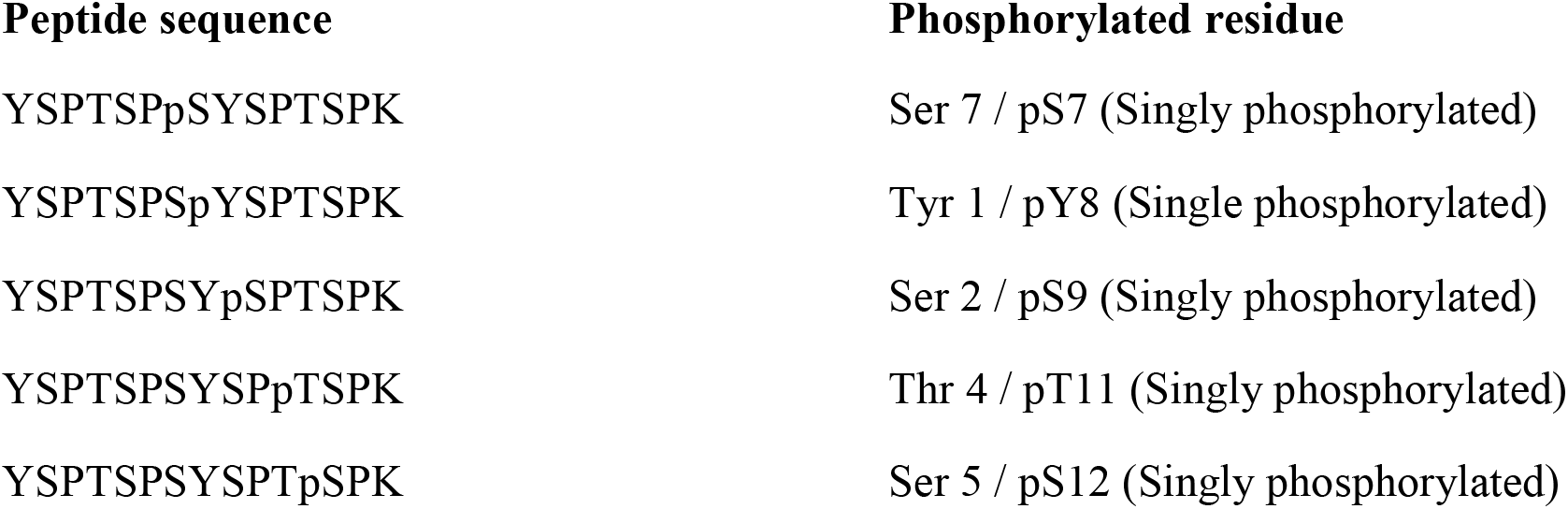
List of diheptad synthetic phosphorylated peptides utilized in this work. Three letter amino acid abbreviations (ex. Ser 5) are used to describe heptad positions, whereas single letters (ex. pS12) designate which residue is phosphorylated within the peptide.

Peptides were analyzed with a benchtop quaternary LC system equipped with a UV detector. To ensure accurate identification of each isoform, we first injected each peptide individually (**Supplementary Figure 2**). Under standard reverse phase conditions (room temp. and pH=2.0) we found that pY8 was readily resolved from the pS/pT peptides, all of which co-eluted regardless of gradient length (**Figure 2A**). The reduced retention and consistent separation of pY from pS/pT peptides suggests more efficient secondary ionization of the tyrosyl phosphate group. We next tested basic mobile phase conditions (pH=10) as well as elevated temperature (65 °C). At each gradient length (including 240 min., **Supplementary Figure 3**) these parameters modestly improved separation, with consistent co-elution of pS9/pT11 and pS7/pS12 (**Figure 2B-D**).

**Figure 2.**
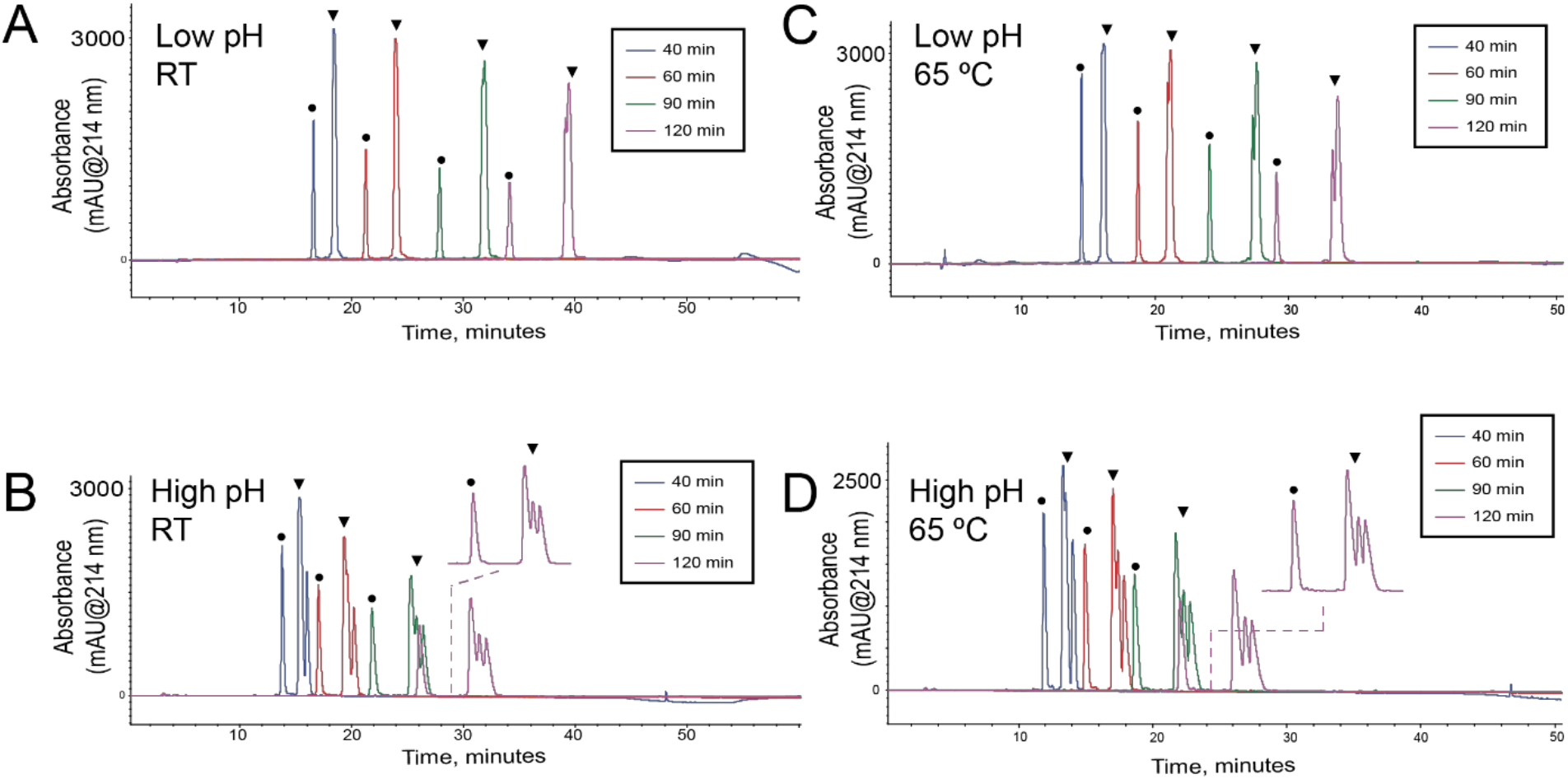
CTD diheptad phosphoisoforms are not well resolved by reversed phase chromatography. A mixture of 10 nmol of each of 5 CTD diheptad phosphoisoforms (Table 1) was analyzed by reversed phase HPLC with gradients ranging from 40 to 120 minutes. (A) pH 2.0, 25 °C. (B) pH 10.0, 25°C. (C) pH 2.0, 65 °C. (D) pH 10.0 65°C. Gradients were 2-40% B in indicated time (A=0.1% TFA,B=acetonitrile with 0.1% TFA). ●, phosphotyrosine peptide;▾ phosphoserine/threonine peptides

**Figure 3.**
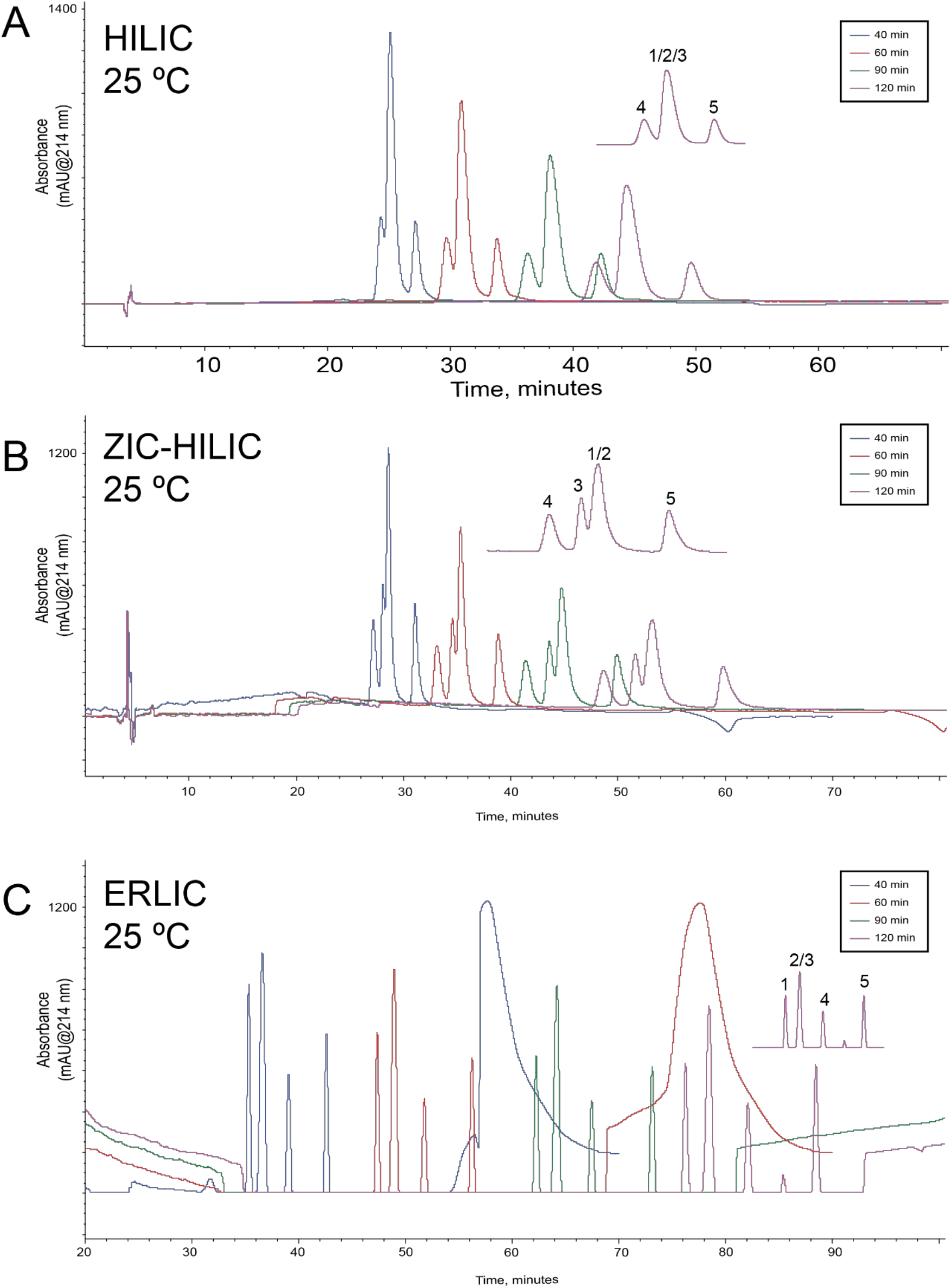
Separation of 10 nmol each of 5 diheptad phosphoisoforms (Table 1) by HILIC (A), ZIC-HILIC (B), and ERLIC (C) chromatography with indicated gradient lengths at 25 °C. For A, gradients were 90-60% B in indicated time (A=0.1% TFA, B=70% acetonitrile/30% isopropyl alcohol/0.1% TFA). For B, gradients were 94.7-63.2% B in indicated time (A=0.05% formic acid, 1 mM ammonium acetate, B=67% acetonitrile, 28.6% isopropyl alcohol, 0.05% formic acid, 1 mM ammonium acetate). For C, gradients were 94.7-21% B in indicated time (A=0.1% formic acid, B=95% acetonitrile, 5% water, 10 mM ammonium acetate). Flow rates were 0.8 mL/min for all experiments. 1. YSPTSPSYSPpTSPK (pT11), 2. YSPTSPpSYSPTSPK (pS7), 3. YSPTSPSYp-SPTSPK (pS9), 4. YSPTSPSYSPTpSPK (pS12), 5. YSPTSPSpYSPTSPK (pY8)

These results led us to consider alternatives to reversed phase chromatography for improved resolution. Several reports have shown that HILIC and ERLIC provide orthogonal separation compared to reversed phase and have been used to separate phosphopeptides [43-50]. To assess whether these techniques could improve resolution of CTD phosphoisomers, we subjected our set of synthetic peptides, individually and as mixtures, to HILIC, ZIC-HILIC, and ERLIC chromatography (**Figure 3** and **Supplementary Figure 4**). HILIC chromatography produced three peaks when the mixture of five peptides was injected (**Figure 3A**). Individual injections established that pS12 eluted first, followed by a co-eluting peak of pS7/pS9/pT11, and then pY8 (**Supplementary Figure 4A**). The reversal in elution order of the tyrosine phosphorylated peptide relative to the pS/pT peptides in HIILC vs. RP chromatography reflects the increased retention of HILIC for hydrophilic analytes. These results confirmed the robust capacity of the stationary phases for separating tyr- and ser-/thr-phosphorylated species. Separation at 65 °C did not significantly impact resolution by HILIC (**Supplementary Figure 4B**).

**Figure 4.**
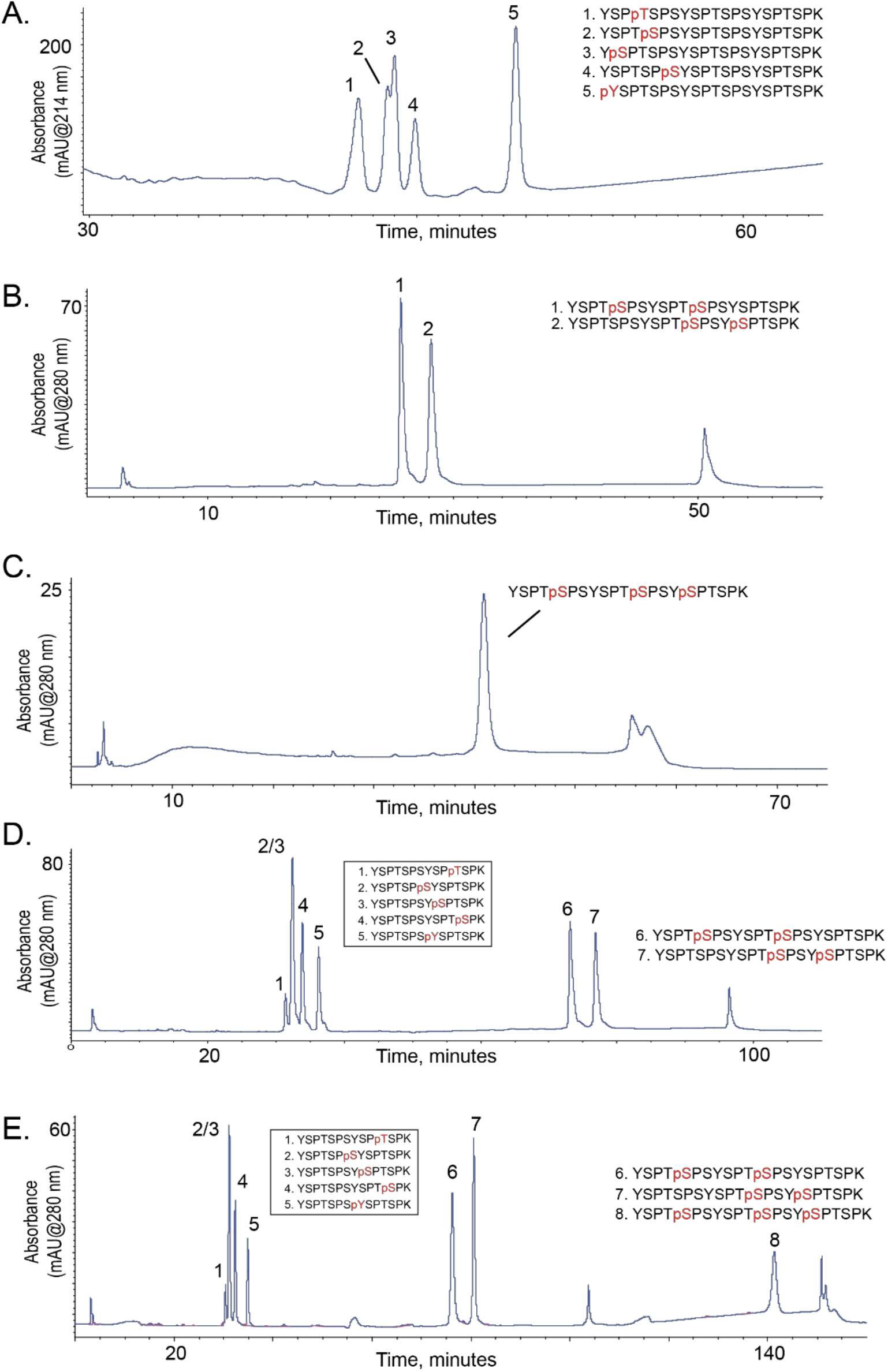
Separation of singly, doubly and triply phosphorylated peptides by ERLIC. (A) Same ERLIC conditions as Figure 3C (60 minute gradient) resolve singly phosphorylated triheptad CTD peptides (2 nmol each injected). (B) 0.2M formic acid elutes doubly phosphorylated triheptad peptides (Table 2) by ERLIC. Gradient was 94.7-5% B in 29 minutes (A=0.2M formic acid, B=95% acetonitrile, 5% water, 0.1% ammonium acetate) and 10 nmol of each peptide was injected. (C) Stronger elution conditions (0.8M formic acid, 50 mM triethylamine) are required to elute triply phosphorylated peptides. Gradient was 94.7-0% B in 34 minutes (A=0.8M formic acid, 50 mM TEA, B=95% acetonitrile, 0.8M formic acid) and 10 nmol of each peptide was injected. (D) A ternary gradient combining conditions of (A) and (B) can resolve singly and doubly phosphorylated peptides in a single analysis. Detailed gradient conditions are described in Supplementary Methods. (E) A quarternary gradient of the solvents in (D) with 0.8M formic acid and 50 mM TEA can separate single, doubly and triply phosphorylated peptides in a single analysis. Detailed gradient conditions are described in Supplementary Methods.

ZIC-HILIC and ERLIC were both superior to reversed phase and HILIC, with ERLIC resolving the mixture of five phosphopeptides clearly into four peaks. Interestingly, pT11 and pS7 co-eluted by ZIC HILIC, whereas pS7 and pS9 coeluted by ERLIC, revealing subtle differences in separation phase selectivity (**Supplementary Figure 4C, 4E**). Elevated temperature improved ZIC HILIC resolution of pS9 from the co-eluting pS7/pT11 peak (**Supplementary Figure 4D**). ERLIC provided improved resolution compared to ZIC-HILIC, even at the shortest gradient tested (40 minutes). Longer gradients at elevated temperature did not improve resolution for HILIC or ZIC-HILIC (**Supplementary Figure 4G, 4H**). The best separation was achieved at elevated temperature with a 240 minute gradient in ERLIC mode, where pS7 and pS9 were partially resolved (**Supplementary Figure 4I**). Although none of the methods surveyed showed complete resolution of all 5 peptides under the conditions tested, ERLIC was clearly superior compared to the other phases tested.

**Table 2.**
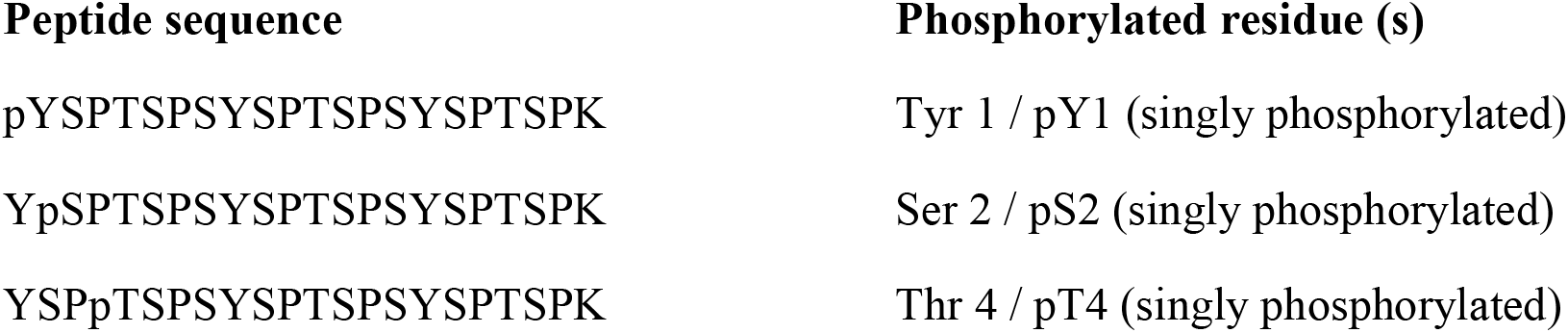
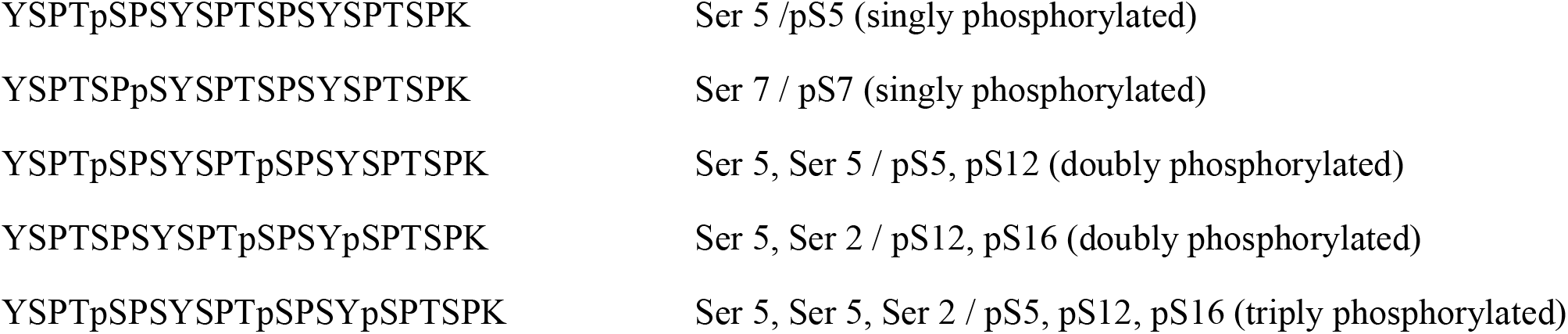
List of triheptad synthetic phosphorylated peptides used in this work. Three letter amino acid abbreviations (ex. Ser 5) are used to describe heptad positions, whereas single letters (ex. pS12) designate which residue is phosphorylated within the peptide.

We next assessed whether ERLIC could provide similar performance for triheptad CTD phosphopeptides. To assess the ability of ERLIC to resolve triheptad peptides, we analyzed several synthetic triheptad phosphoisoforms (**Table 2, Figure 4**). A mixture of five triheptad, singly phosphorylated CTD phosphopeptides was resolved into four peaks (**Figure 4A**), very similar to results with diheptad isoforms. Taken together, these results highlight the improved ability of ERLIC to resolve multiple groups of related CTD phosphoisoforms.

To determine compatibility of ERLIC with the analysis of multiply phosphorylated CTD peptides, we first analyzed a mixture of two doubly phosphorylated triheptad peptides by ERLIC using the same gradient conditions but found that the peptides did not elute (data not shown). This result is consistent with the strong retention of the phosphate group on the ERLIC column. Work by Cui *et al*. demonstrated that stronger organic modifiers such as triethylamine phosphate or TFA can mediate elution of multiply-phosphorylated peptides from ERLIC [48]. We sought to avoid salts or TFA to maintain compatibility with online elution of ERLIC directly to a mass spectrometer for sequence identification. We reasoned that higher concentrations of formic acid may reduce retention via protonation of the phosphate group, while maintaining sufficient electrospray ionization. Indeed, increasing the concentration of formic acid in the A solvent to 0.2M (from 0.1%) led to elution of resolved, doubly phosphorylated peptides (**Figure 4B**). However, elution of the triply-phosphorylated peptide required 0.8M formic acid and 50 mM triethylamine (**Figure 4C**). Interestingly, by combining elution conditions in ternary or quaternary gradients, we achieved separation of singly and doubly (**Figure 4D**) or singly, doubly and triply (**Figure 4E**) phosphorylated peptides by ERLIC.

We next sought to leverage these conditions for the analysis of msCTD phosphopeptides by capillary-scale LC-MS/MS. For these studies we utilized a 75 µm × 15 cm ERLIC column with an integrated emitter tip operated at 400 nL/min interfaced to a QTOF mass spectrometer[51]. Mixtures of five di- and tri-heptad msCTD phosphopeptides were then separated using a gradient from 94.7%-40% B in 40 minutes (A=0.1% formic acid, B=0.01% ammonium acetate in 95% acetonitrile). Ion chromatograms (**Figure 5A,B**) were very similar to UV chromatograms generated with the 4.6 mm ERLIC column. The five diheptad phosphopeptides were well resolved into four peaks with pS7/pS9 co-eluting in the second peak (**Figure 5A**). All five triheptad phosphopeptides were observed as distinct peaks, although not baseline resolved (**Figure 5B**). We next replaced solvent A with 0.2M formic acid and analyzed a mixture of two doubly phosphorylated triheptad phosphopeptides using the same gradient. As illustrated in **Figure 5C** both peptides were resolved and detected. These elution conditions could be utilized with a ternary gradient to elute and resolve both types of peptides in a single analysis.

**Figure 5.**
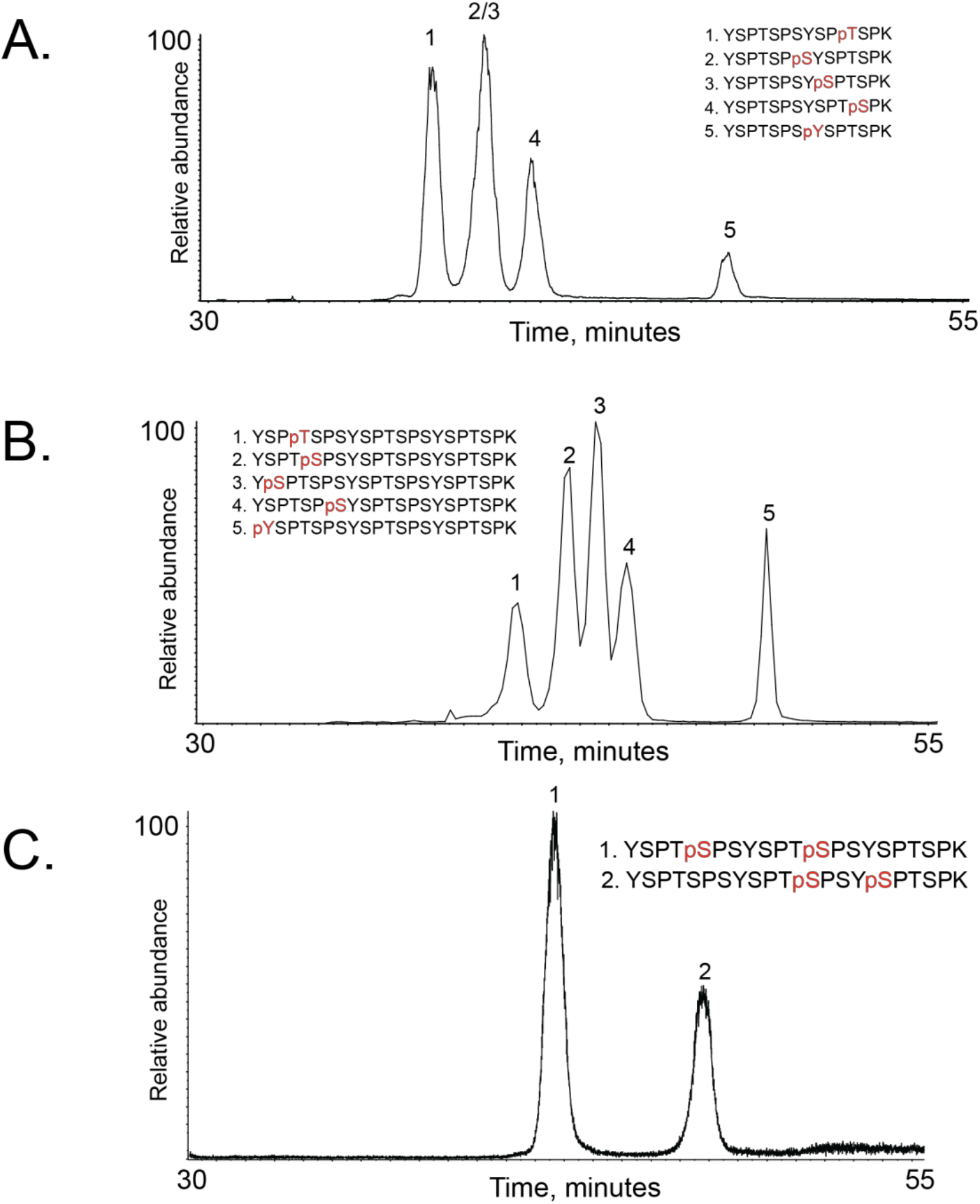
Nanoscale ERLIC-MS analysis of singly phosphorylated diheptad (A), singly phosphorylated triheptad (B), and doubly phosphorylated triheptad (C) phosphopeptides. Detailed gradient conditions are given in Supplementary Methods.

In preparation for LC-MS/MS analysis of triply phosphorylated msCTD peptides with TEA as the organic modifier, we infused a singly phosphorylated synthetic peptide (CTD pS5) in a solution of 30% acetonitrile with either 0.1% formic acid, or 0.8M formic acid with 50 mM triethylamine (as a mimic of ERLIC gradient conditions). Unfortunately, use of TEA led to significant ionization suppression, suggesting incompatibility with online LC-MS/MS acquisition (data not shown). This result suggests that CTD peptides with higher-order phosphorylation should be analyzed by ZIC-HILIC, albeit with somewhat reduced resolution (**Supplementary Figure 5**).

We next sought to evaluate the ERLIC approach in the context of full length CTD modified by bona fide CTD kinases. Our earlier study using native msCTD found no obvious differences in phosphorylation at specific repeats positioned throughout the CTD [39]. We therefore created a second generation msCTD with lysine residues present at either every second (msCTD2.2K) or third heptad repeat (msCTD2.3K) (**Figure 1B**). While these versions lack encoding to distinguish repeat position, they provide an amplification of lower abundance phosphoisomers by summing signals across repeat sequences throughout the CTD.

We first generated recombinant msCTD2.2K as a GST fusion. Purified protein was phosphorylated with the Ser5 kinase Kin28 (known as Cdk7 in metazoans) that was isolated from yeast using tandem affinity purification as previously described [39]. Strong phosphorylation was confirmed by immunoblotting (**Figure 6A**). We then digested the phosphorylated msCTD2.2K with trypsin, and analyzed peptides with online ERLIC-MS/MS using targeted MS^2^ scans to detect doubly charged, single phosphorylated diheptad peptides. MS/MS spectra (**Supplementary Figure 6**) demonstrate complete resolution of Ser5 phosphorylation in first and second heptad repeats (1. Ser 5 / pS5 and 3. Ser 5 / pS12 in **Figure 6B**). Notably, we also detect a small signal (∼30 fold less abundant based on peak area) corresponding to Ser 2 phosphorylation in the second heptad (2. Ser 2 / pS9), confirming that on-line ERLIC MS/MS can resolve and detect phosphoisoforms that co-elute by RP chromatography (**Figure 2**).

**Figure 6.**
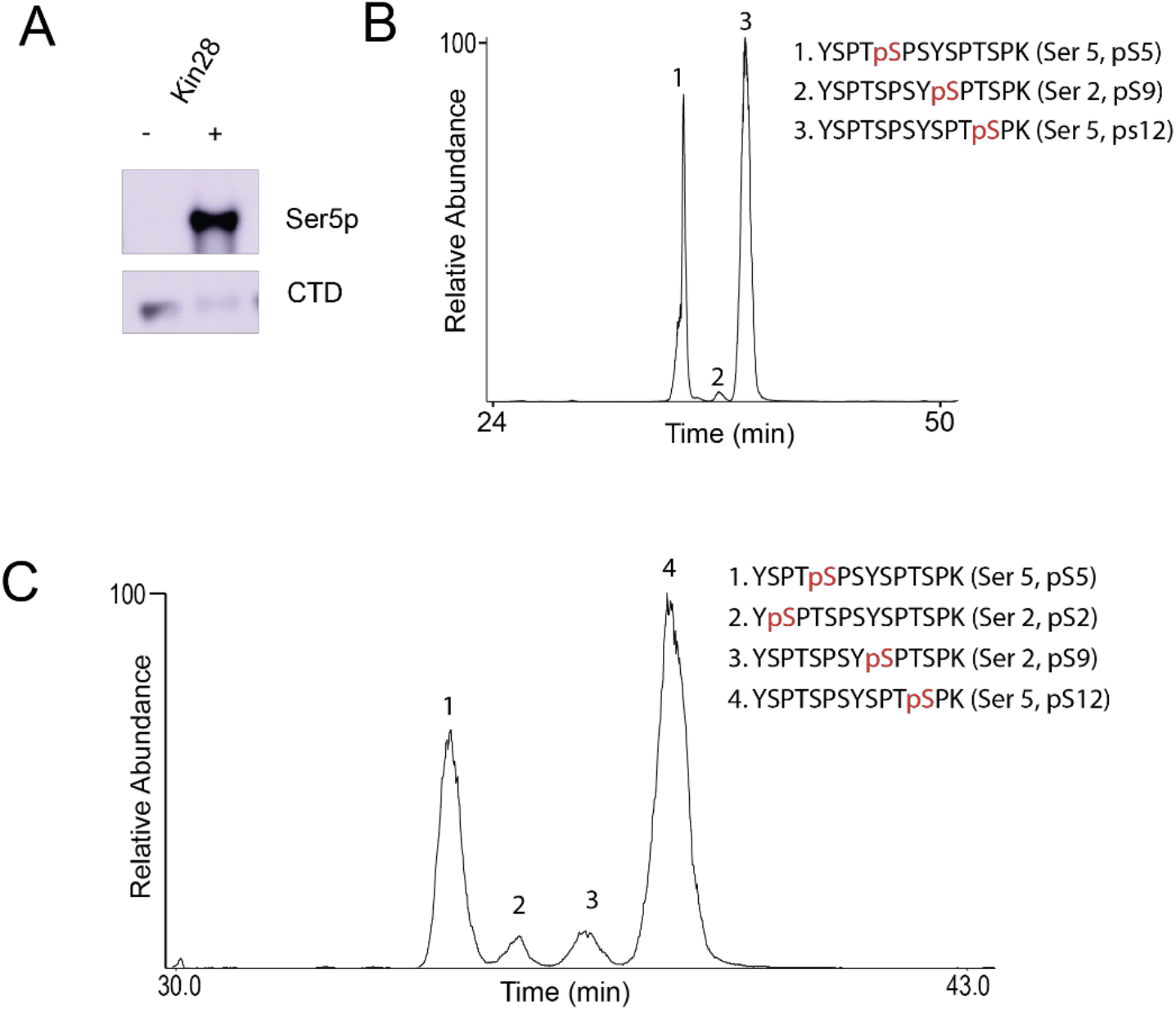
On-line ERLIC analysis of biologically derived CTD phosphopeptides. (A) Immunoblot of recombinant msCTD2.2K after in vitro phosphorylation with the Ser5 kinase Kin28. (B, C) Extracted ion chromatogram of doubly charged, singly phosphorylated diheptad CTD peptide YSPTSPSYSPTSPK acquired during analysis of (B) Kin28 treated msCTD2.2K tryptic peptides, or (C) msCTD2.2K isolated from native RNA pol II expressed in yeast cells. Gradient conditions were the same as Figure 5A (see Supplementary Methods).

We next purified bulk RNA pol II from yeast cells expressing msCTD2.2K as part of the sole Rpb1 gene. The msCTD2.2K fragment was purified as previously described [39], subjected to trypsin digestion, and the resulting phosphopeptides were analyzed by online ERLIC MS/MS using the same targeted MS^2^ method (**Figure 6C**). In this experiment, we observed resolution of Ser 2 (pS2 / pS9) and Ser5 (pS5 / pS12) phosphopeptides within the first and second heptads. Confirming our earlier analysis, Ser5 phosphopeptides were more abundant than those of Ser2 (∼10 fold based on peak area). These results strongly validate ERLIC as a superior method for resolving phosphopeptide isoforms within the C-terminal domain of RNA polymerase II.

## CONCLUSIONS AND OUTLOOK

We performed an in-depth study of how synthetic CTD peptides resolve by reversed phase, HILIC, ZIC-HILIC, and ERLIC chromatography under a variety of conditions. We found that standard reversed phase chromatography is insufficient to resolve a majority of serine/threonine phosphoisoforms, even at elevated temperatures and with extended gradients. In contrast, ZIC-HILIC and ERLIC offer superior resolution of phosphopeptide isoforms and furthermore we identified buffer conditions that are compatible with online sequence analysis by mass spectrometry. Our progress herein sets the stage for future studies focused on quantification of individual CTD phosphorylation sites. For example, incorporation of isotopically encoded (AQUA) CTD phosphopepitde analogs in our ERLIC workflow could be used to establish not only absolute abundance levels but also site occupancy of resolved isoforms[52]. Also, it is interesting to note that our analysis of RNA pol II from yeast cells expressing msCTD2.2K detected no Tyr 1, Thr 4, or Ser7 phosphorylation. Assuming roughly similar ionization efficiency of the phospho-isoforms, this observation suggests they are present at far lower abundance than Ser 5 and Ser 2. In future experiments we can use our ERLIC platform to interrogate msCTD phosphorylation with high temporal resolution at different stages of transcription to better delineate dynamics and relative contribution of transient modifications.

Beyond the systematic evaluation reported herein, our future work will focus on identifying buffer conditions for separation and mass spectrometry analysis of triply phosphorylated CTD peptides. Notwithstanding this limitation, our results demonstrate the potential for ERLIC to facilitate analysis of *in vivo* CTD phosphorylation to improve our understanding of how multiple signaling axes regulate gene transcription. More generally, efforts in functional proteomics to annotate the dynamics of phosphorylation at proximal residues on protein substrates will benefit tremendously from further advances in ERLIC separation platforms.

## Supporting Information

Electronic supplementary information is available at the publisher’s website as a single pdf containing Supplementary Figures and Supplementary Methods.

The Supporting Information is available free of charge on the ACS Publications website.

## ACKNOWLEDGMENT

The authors acknowledge Andy Alpert for his generous commitment of time and insightful guidance for HILIC and ERLIC methods in support of this project. This work was supported by NIH award GM056663. We are grateful to Natalia Orlovsky for technical assistance with second generation msCTD work, and to all members of the Marto and Buratowski labs for helpful discussions.

## CRediT authorship contribution statement

**Scott Ficarro:** Investigation, Methodology, Visualization, Formal Analysis, Supervision, Writing – original draft, review and editing. **Deepash Kothiwal:** Investigation, Methodology, Visualization, Formal Analysis, Writing – original draft, review and editing. **Hyun Jin Bae:** Investigation, Methodology, Visualization. **Isidoro Tavares:** Investigation. **Gabriela Giordano:** Investigation. **Stephen Buratowski:** Funding Acquisition, Supervision, Visualization, Resources, Writing – review and editing. **Jarrod A. Marto:** Conceptualization, Funding Acquisition, Supervision, Visualization, Resources, Writing – original draft, review and editing.

## Supplementary Figure Legends

**Supplementary Figure 1**. (A) Extracted ion chromatogram for m/z 789.84 corresponding to singly phosphorylated, doubly charged YSPTSPSYSPTSPK acquired during analysis of msCTD2.2K peptides derived from yeast. The split, poorly resolved peak is indicative of multiple, co-eluting phosphoisoforms. (B) MS/MS spectrum of singly phosphorylated, doubly charged YSPTSPSYSPTSPK peptide obtained at 41.07 minutes (marked with red ‘*’ in A) shows fragment ions consistent with phosphorylation of both the first (▲) and second (*) heptads.

**Supplementary Figure 2**. UV traces of 10 nmol diheptad phosphoisoform peptides S7, Y8, S9, T11, and S12 by RP chromatography at (A) low pH [A=0.1% TFA in water,B=0.1% TFA in acetonitrile] or (B) high pH [A=20 mM ammonium formate in water, B=acetonitrile). Gradient was 2-40% B in 120 minutes with a flow rate of 0.8 mL/min at 65 °C with UV detection at 214 nm.

**Supplementary Figure 3**. UV traces of a mixture of 10 nmol each of diheptad phosphoisoform peptides S7, Y8, S9, T11, and S12 by RP chromatography at (A) low pH [A=0.1% TFA in water,B=0.1% TFA in acetonitrile] or (B) high pH [A=20 mM ammonium formate in water, B=acetonitrile). Gradient was 2-40% B in 240 minutes with a flow rate of 0.8 mL/min at 65 °C with UV detection at 214 nm.

**Supplementary Figure 4**. (A) UV traces of 10 nmol diheptad phosphoisoform peptides S7, Y8, S9, T11, and S12 by HILIC chromatography. Gradient was 90-60% B in 120 minutes (A=0.1% TFA, B=70% acetonitrile/30% isopropyl alcohol/0.1% TFA) (B) UV traces of a mixture of 10 nmol each of diheptad phosphoisoform peptides S7, Y8, S9, T11, and S12 by HILIC chromatography with gradients of 90-60% B in 40, 60, 90, or 120 minutes (A=0.1% TFA, B=70% acetonitrile/30% isopropyl alcohol/0.1% TFA) (C) UV traces of 10 nmol diheptad phosphoisoform peptides S7, Y8, S9, T11, and S12 by ZIC-HILIC chromatography. Gradient was 94.7-63.2% B in 120 minutes (A=0.05% formic acid, 1 mM ammonium acetate, B=67% acetonitrile/28.6% isopropyl alcohol/0.05% formic acid, 1 mM ammonium acetate) (D) UV traces of a mixture of 10 nmol each of diheptad phosphoisoform peptides S7, Y8, S9, T11, and S12 by ZIC-HILIC chromatography with gradients of 94.7-63.2% B in 40, 60, 90, or 120 minutes (A=0.05% formic acid, 1 mM ammonium acetate, B=67% acetonitrile/28.6% isopropyl alcohol/0.05% formic acid, 1 mM ammonium acetate) (E) UV traces of 10 nmol diheptad phosphoisoform peptides S7, Y8, S9, T11, and S12 by ERLIC chromatography. Gradient was 94.7-21% B in 40 minutes (A=0.1% formic acid, B=95% acetonitrile with 10 mM ammonium acetate) (F) UV traces of a mixture of 10 nmol each of diheptad phosphoisoform peptides S7, Y8, S9, T11, and S12 by ERLIC chromatography with gradients of 94.7-21% B in 40, 60, 90, or 120 minutes (A=0.1% formic acid, B=95% acetonitrile with 10 mM ammonium acetate) (G-I) UV traces of a mixture of 10 nmol each of diheptad phosphoisoform peptides S7, Y8, S9, T11, and S12 by (G) HILIC, (H) ZIC-HILIC, and (I) ERLIC chromatography. (G) Gradient was 90-60% B in 240 minutes (A=0.1% TFA, B=70% acetonitrile/30% isopropyl alcohol/0.1% TFA) (H) Gradient was 94.7-63.2% B in 240 minutes (A=0.05% formic acid, 1 mM ammonium acetate, B=67% acetonitrile/28.6% isopropyl alcohol/0.05% formic acid, 1 mM ammonium acetate) (I) Gradient was 94.7-21% B in 240 minutes (A=0.1% formic acid, B=95% acetonitrile with 10 mM ammonium acetate). All experiments used a flow rate of 0.8 mL/min, a column temperature of 65 °C and UV detection at 214 nm. (B,D,F-I) 1. YSPTSPSYSPpTSPK (pT11), 2. YSPTSPpSYSPTSPK (pS7), 3. YSPTSPSYpSPTSPK (pS9), 4. YSPTSPSYSPTpSPK (pS12), 5. YSPTSPSpYSPTSPK (pY8)

**Supplementary Figure 5**. Triply phosphorylated CTD peptides can be eluted in mass spectrometry compatible solvents using ZIC-HILIC by offline (A) or online to the mass spectrometer (B). For (A), the gradient was 90-60% B in 60 minutes, A=1 mM ammonium acetate in water/0.05% formic acid, B=67% acetonitrile, 1 mM ammonium acetate, 0.05% formic acid, 28.6% isopropyl alcohol, flow rate 0.8 mL/min, 65 °C. For (B), the gradient was 90-50% B in 5 minutes, A=1 mM ammonium acetate in water/0.05% formic acid, B=67% acetonitrile, 1 mM ammonium acetate, 0.05% formic acid, 28.6% isopropyl alcohol, flow rate 400 nL/min, room temperature, with 400 fmol injected.

**Supplementary Figure 6**. MS/MS spectra of phosphopeptides acquired during ERLIC-MS analysis of Kin28 treated msCTD2.2K. (A) MS/MS spectrum of YSPTpSPSYSPTSPK (Ser 5, pS5). Ions y9, y10, and y10Δ indicate phosphorylation of pS5. (B) MS/MS spectrum of YSPTSPSYpSPTSPK (Ser 2, pS9). Ions y5, y6, and y6Δ indicate phosphorylation of pS9. (C) MS/MS spectrum of YSPTSPSYSPTpSPK (Ser 5, ps12). Ions y2, y3, and y3Δ indicate phosphorylation of pS12. Δ, loss of H3PO4.

**Supplementary Figure 7**. MS/MS spectra of phosphopeptides acquired during ERLIC-MS analysis of msCTD2.2K from yeast cells. (A) MS/MS spectrum of YSPTpSPSYSPTSPK (Ser 5, pS5). Ions y9, y10, and y10Δ indicate phosphorylation of pS5. (B) MS/MS spectrum of YpSPTSPSYSPTSPK (Ser 2, pS2). Ion y12 places the phosphate on pY1 or pS2. Neutral loss of phosphate (absent from pY peptides) as well as elution before pS9/pS12 peptides indicate pS2 is the phosphorylated residue. (C) MS/MS spectrum of YSPTSPSYpSPTSPK (Ser 2, pS9). Ions y5, y6, and y6Δ indicate phosphorylation of pS9. (D) MS/MS spectrum of YSPTSPSYSPTpSPK (Ser 5, pS12). Ions y2, y3, and y3Δ indicate phosphorylation of pS12. Δ, loss of H3PO4.

