## Supplemental Figures and Methods for "Leveraging HILIC/ERLIC Separations for Online Nanoscale LC-MS/MS Analysis of Phosphopeptide Isoforms from RNA Polymerase II C-terminal Domain"

### Supplementary Methods

##### Detailed gradient conditions for Figure 4

**Figure 4D.**

Buffer A: 2%MeCN/0.1% formic acid

Buffer B: 95% Acetonitrile with 0.1% ammonium acetate, 5% water

Buffer C: 0.2M formic acid

| Time | A% | B% | C% |
|---|---|---|---|
| 0 | 5.3 | 94.7 | 0 |
| 1 | 5.3 | 94.7 | 0 |
| 40 | 95 | 5 | 0 |
| 45 | 95 | 5 | 0 |
| 50 | 5.3 | 94.7 | 0 |

| | | | |
|---|---|---|---|
| 51 | 0 | 94.7 | 5.3 |
| 81 | 0 | 5 | 95 |
| 91 | 0 | 5 | 95 |
| 96 | 0 | 94.7 | 5.3 |
| 97 | 5.3 | 94.7 | 0 |
| 102 | 5.3 | 94.7 | 0 |

10 nmol of each peptide was injected in a volume of 50 uL.

**Figure 4E.**

Buffer A: 2%MeCN/0.1% formic acid

Buffer B: 95% Acetonitrile with 0.1% ammonium acetate, 5% water

Buffer C: 0.2M formic acid

Buffer D: 0.8M formic acid, 50mM TEA

| Time | A% | B% | C% | D% |
|---|---|---|---|---|
| 0 | 5.3 | 94.7 | 0 | 0 |
| 1 | 5.3 | 94.7 | 0 | 0 |
| 45 | 95 | 5 | 0 | 0 |
| 50 | 95 | 5 | 0 | 0 |
| 55 | 5.3 | 94.7 | 0 | 0 |
| 56 | 0 | 94.7 | 5.3 | 0 |
| 58 | 0 | 94.7 | 5.3 | 0 |
| 88 | 0 | 5 | 100 | 0 |
| 98 | 0 | 5 | 100 | 0 |
| 103 | 0 | 94.7 | 5.3 | 0 |
| 104 | 0 | 94.7 | 0 | 5.3 |
| 106 | 0 | 94.7 | 0 | 5.3 |
| 136 | 0 | 0 | 0 | 100 |
| 146 | 0 | 0 | 0 | 100 |
| 151 | 0 | 94.7 | 0 | 5.3 |
| 156 | 0 | 94.7 | 0 | 5.3 |

10 nmol of each peptide was injected in a volume of 50 uL.

#### Detailed gradient conditions for Figure 5

(A) Total ion chromatogram for MS/MS scans of the doubly charged, singly phosphorylated diheptad peptides, 375 fmol of each peptide was injected, gradient 94.7-50% B in 40 minutes, flow rate 400 nL/min, A=0.1% formic acid, B=95% acetonitrile with 0.05% ammonium acetate, 5% water. (B) Extracted ion chromatogram for singly phosphorylated triheptad phosphopeptides. 50 fmol of each peptide was injected. Gradient, flow rate, and solvent compositions were the same as in(A). (C) Extracted ion chromatogram for the doubly phosphorylated triheptad phosphopeptides. 200 fmol of each peptide was injected. Gradient was 94.7-30% B in 34 minutes, A=0.2M formic acid, B=95% acetonitrile with 0.05% ammonium acetate, 5% water.

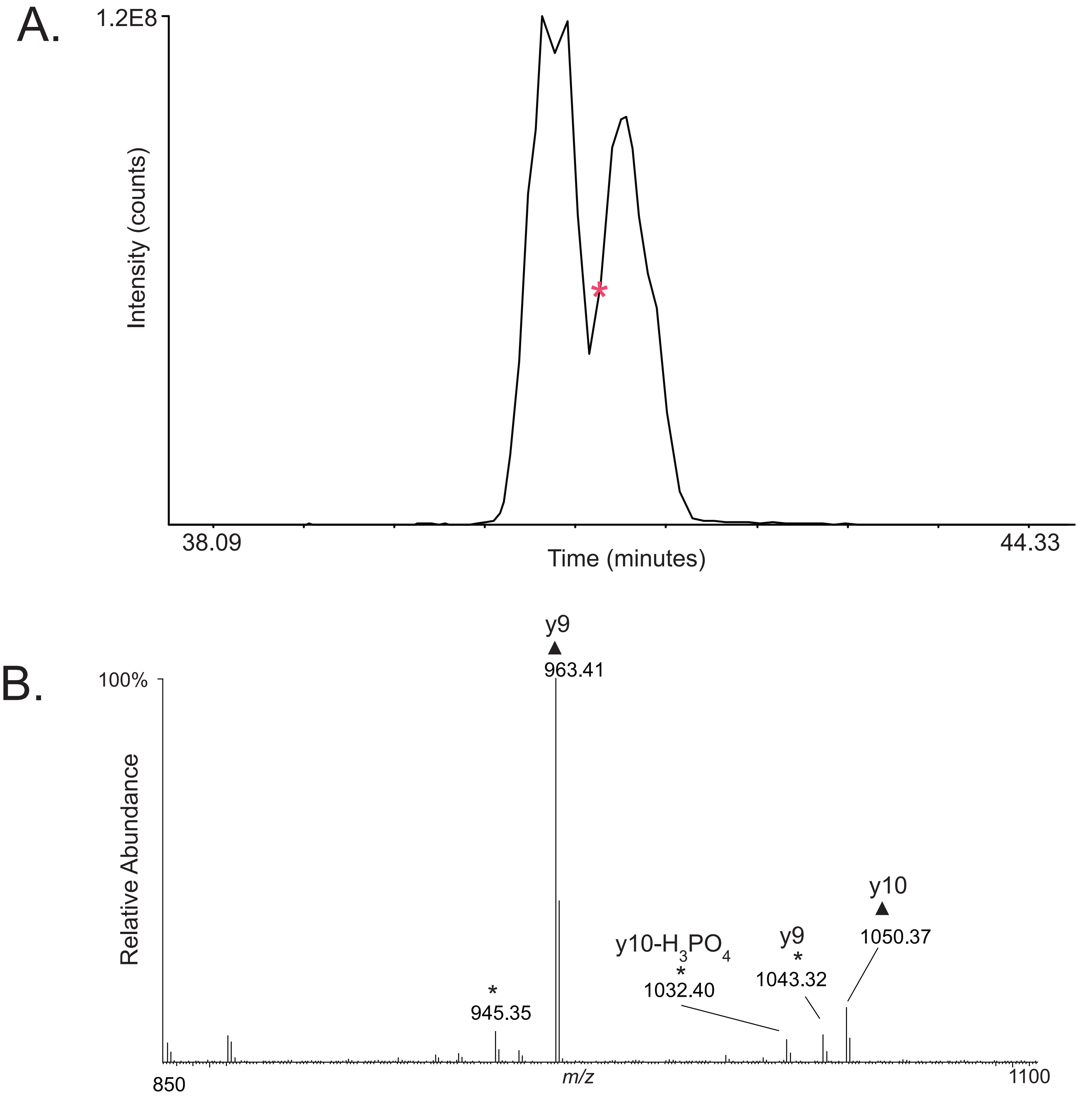


Ficarro et al Supplementary Figure 1.

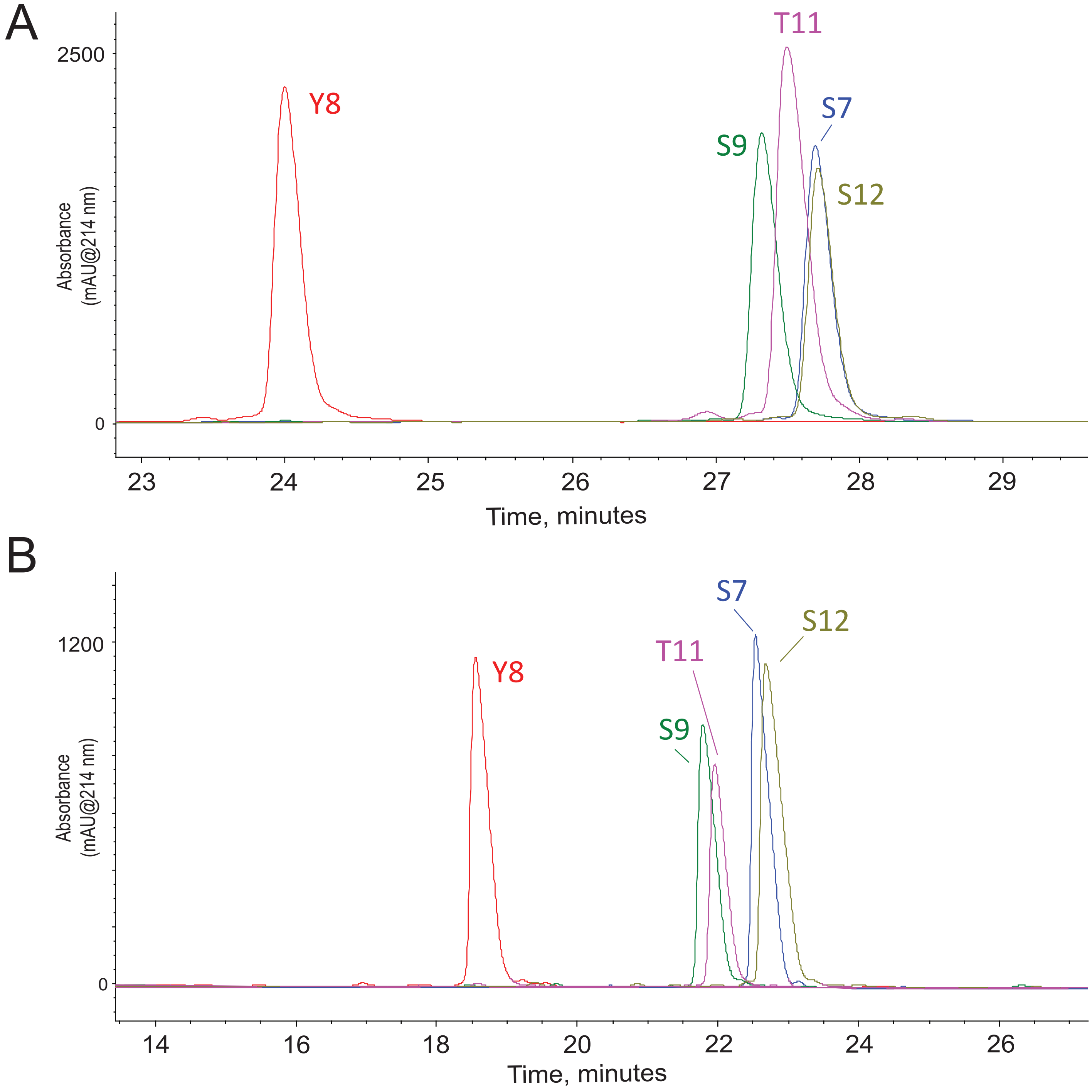


Ficarro et al Supplementary Figure 2.

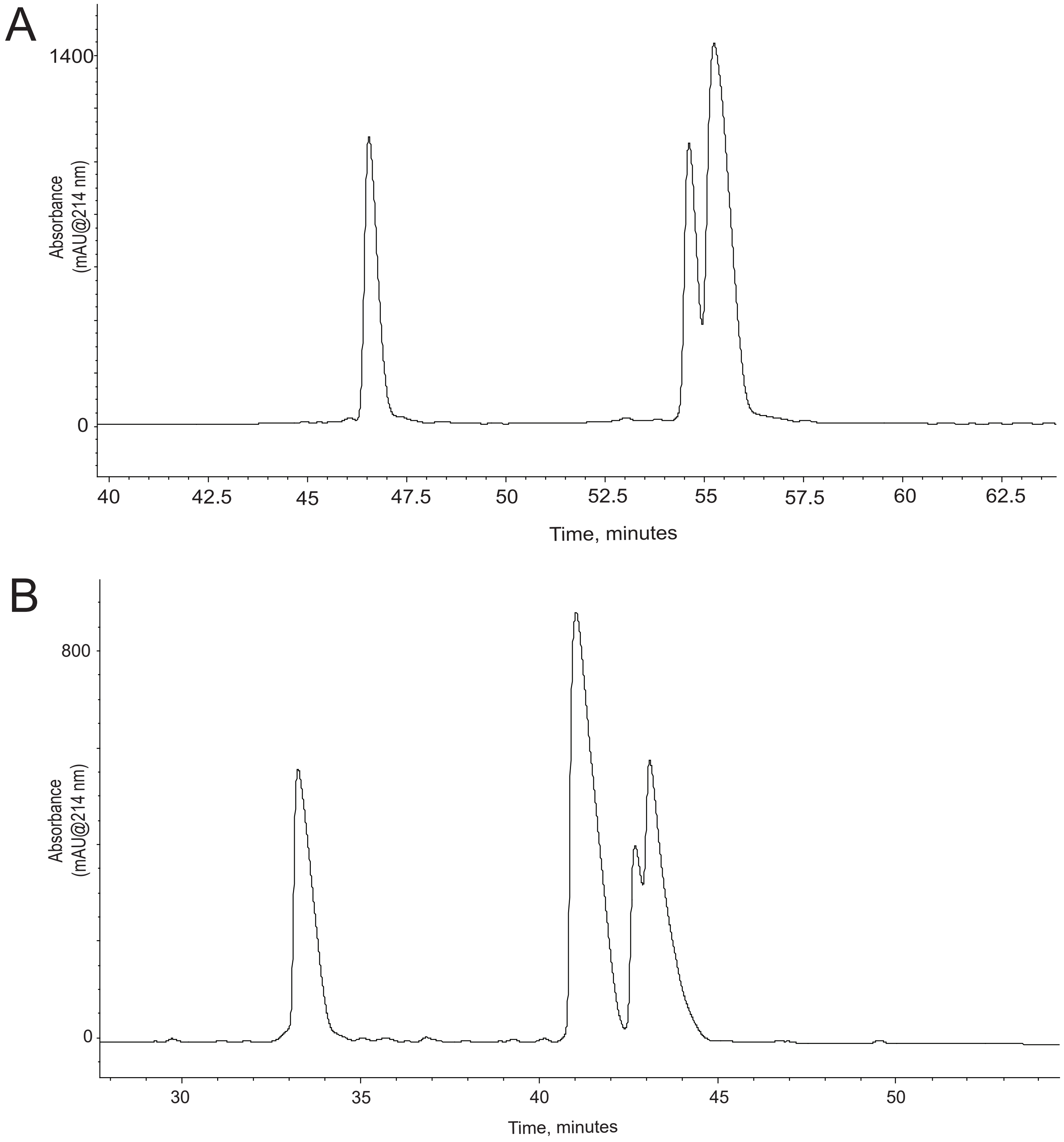


Ficarro et al Supplementary Figure 3.

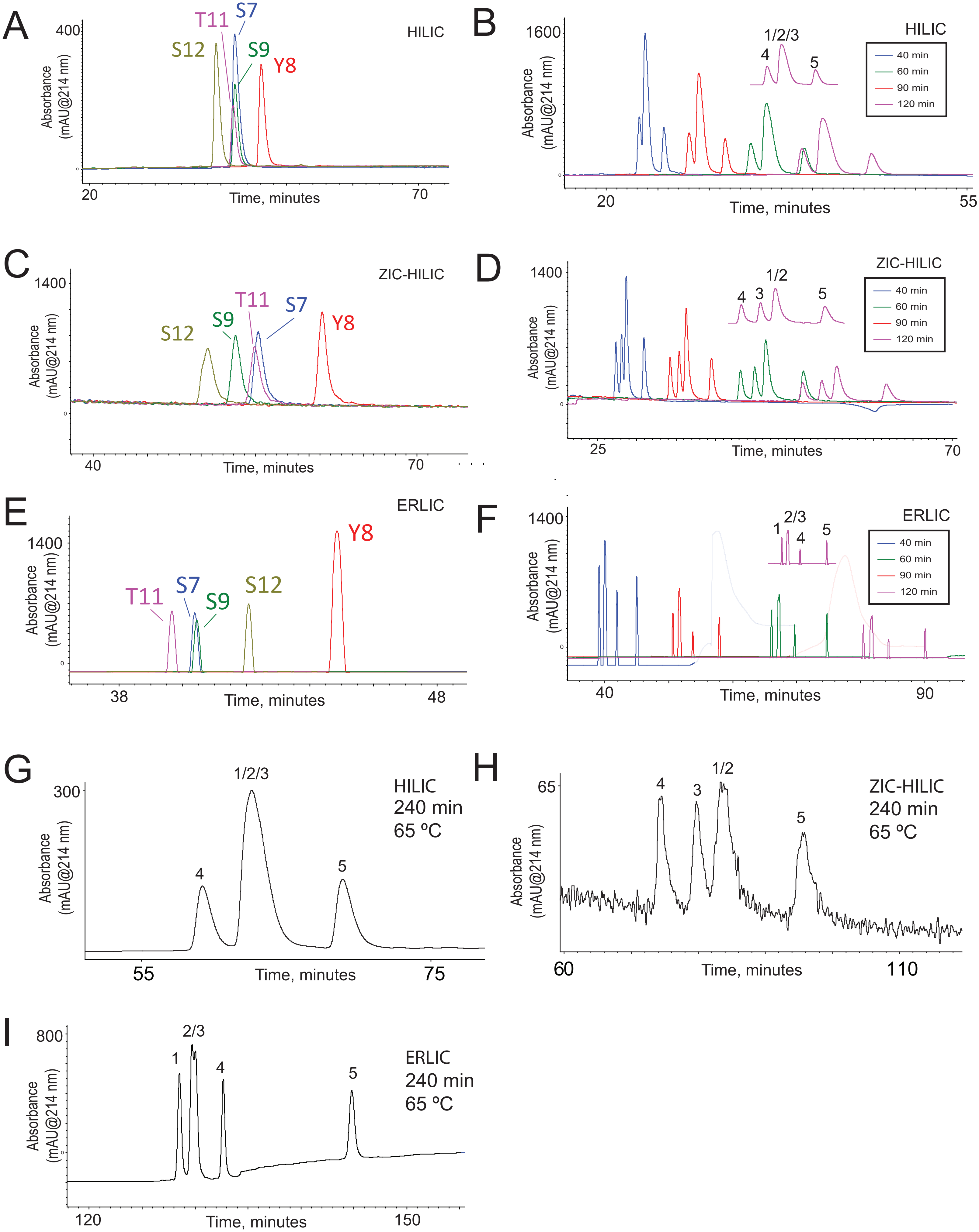


Ficarro et al Supplementary Figure 4.

A.

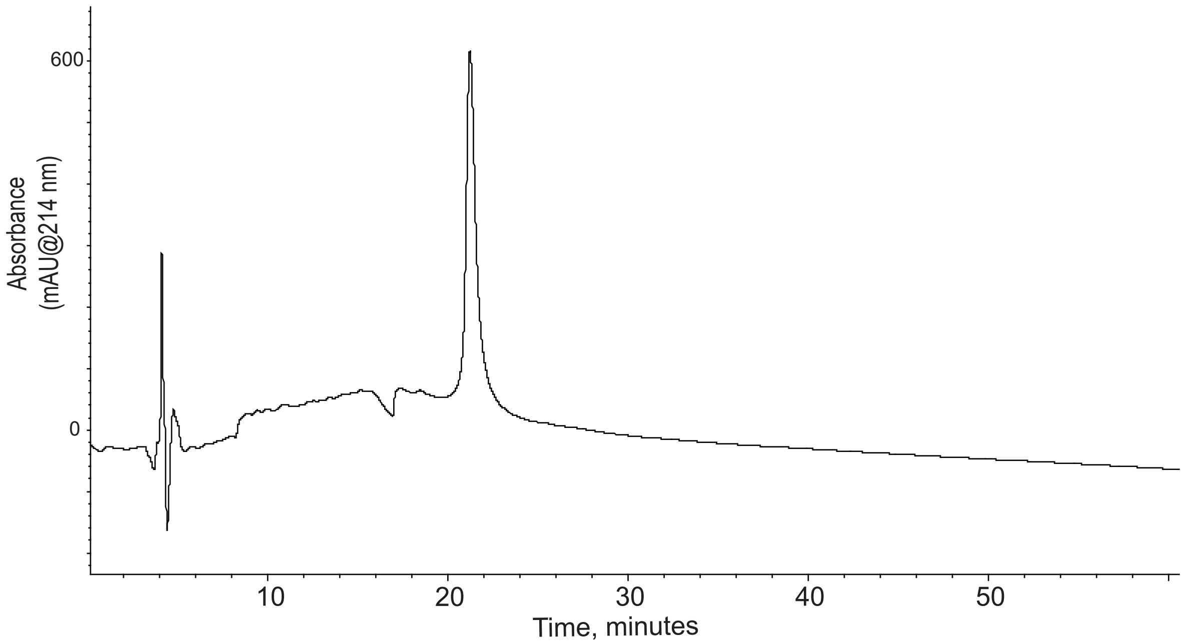


B.

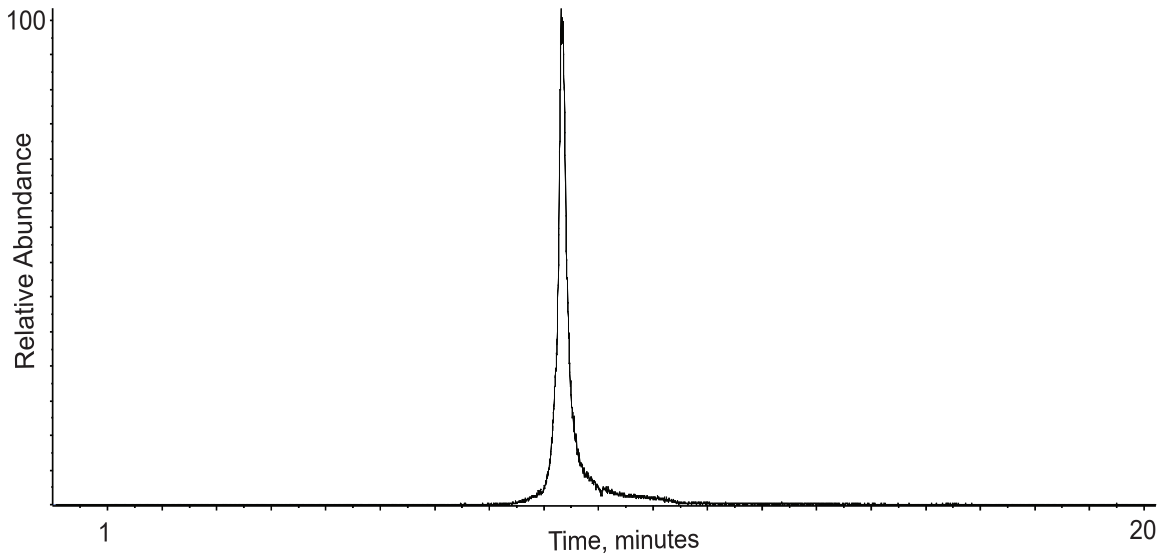


Ficarro et al Supplementary Figure 5.

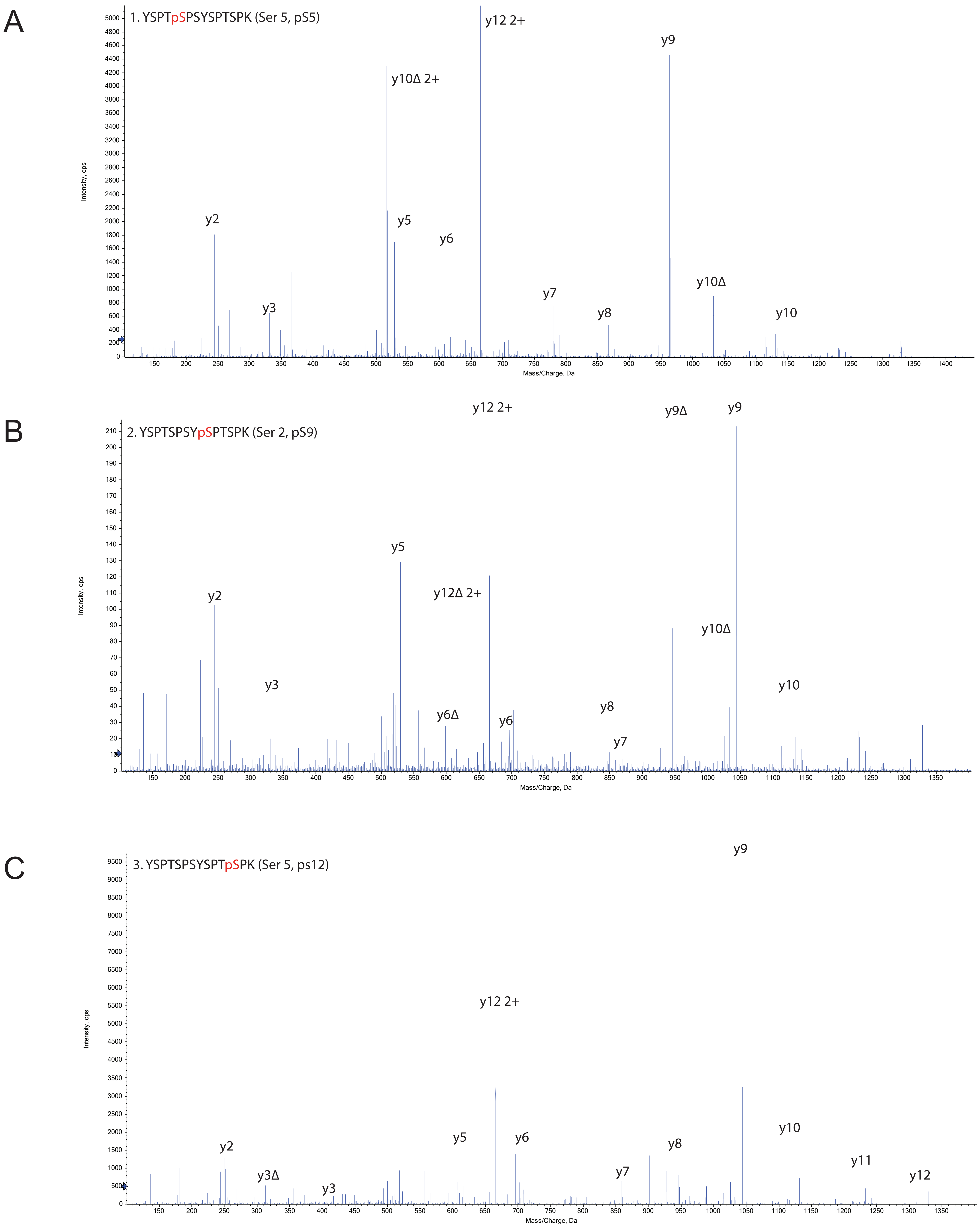


Ficarro et al Supplementary Figure 6.

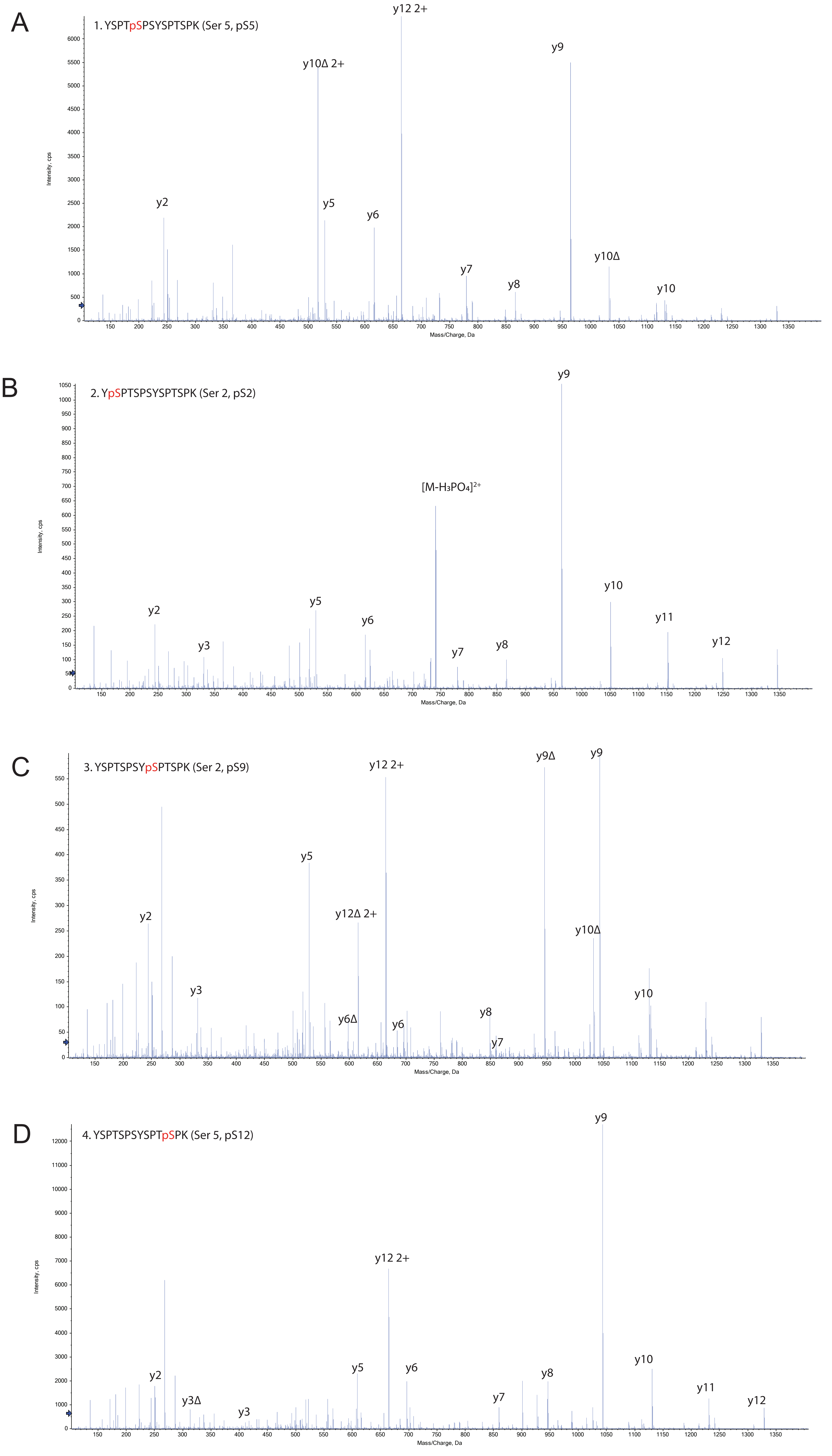


Ficarro et al Supplementary Figure 7.
